## Supplementary Tables 1-2 and Suppl Figures 1-7 for "Stranded short nascent strand sequencing reveals the topology of DNA replication origins in *Trypanosoma brucei*"

^#^ Authors contributed to the manuscript equally

### SUPPLEMENTARY DATA

##### Supplementary Table 1. Numbers of reads

|  |  |  |  | **Peaks called against SNS depleted control** | | | |
| --- | --- | --- | --- | --- | --- | --- | --- |
|  | **Total reads** | **Reads after trimming and deduplication** | **Mapped reads** | **(+) peaks called** | **(-) peaks called** | **filtered peaks (+) and (-)** | **mapped ORIs** |
| PCF repl 1 SNS enriched | 35377234 | 26331533 | 1990221 | 890 | 919 | 76 | 38 |
| PCF repl 1 SNS depleted | 75161248 | 48009990 | 3310426 |  |  |  |  |
| PCF repl 2 SNS enriched | 40265002 | 29772101 | 2119887 | 2101 | 2660 | 170 | 85 |
| PCF repl 2 SNS depleted | 44506300 | 37022757 | 2724281 |  |  |  |  |
| PCF repl 3 SNS enriched | 67335770 | 57786492 | 4206187 | 5788 | 5381 | 904 | 452 |
| PCF repl 3 SNS depleted | 34346800 | 28214856 | 1999576 |  |  |  |  |
| BSF repl 1 SNS enriched | 32609118 | 21305027 | 1289981 | 2028 | 1866 | 462 | 231 |
| BSF repl 1 SNS depleted | 32673418 | 14195198 | 5432187 |  |  |  |  |
| BSF repl 2 SNS enriched | 49794683 | 43417511 | 2917335 | 7592 | 8268 | 1436 | 718 |
| BSF repl 2 SNS depleted | 42316922 | 36253185 | 2682328 |  |  |  |  |
| BSF repl 3 SNS enriched | 28145055 | 22793377 | 1548353 | 2986 | 3158 | 614 | 307 |
| BSF repl 3 SNS depleted | 28234323 | 21022409 | 1411179 |  |  |  |  |

The numbers of reads obtained after paired-end sequencing, numbers of reads, peaks and mapped origins after different steps of bioinformatic analysis are presented for three biological replicates of PCF and BSF cells. Each replicate contained an SNS-enriched sample and SNS-depleted control. (+) and (-) peaks called presents the number of peaks called and localized on the plus or minus DNA strand, respectively. Filtered peaks (+) and (-) represent pears of peaks with divergent orientation (first (-) followed by (+) peak) (Methods).

##### Supplementary Table 2. **DNA replication parameters obtained by DNA combing**

|  |  | **PCF** | **BSF** |
| --- | --- | --- | --- |
|  | **Inter-origin distance (kb)** | | |
|  | Number of values | 144 | 101 |
|  | Minimum | 23.26 | 37.2 |
|  | 25% Percentiles | 106.1 | 145.7 |
|  | Median | 152 | 212.8 |
|  | 75% Percentiles | 227.1 | 284.1 |
|  | Maximum | 597 | 584.9 |
|  | Mean | 179.4 | 219.4 |
|  | **Velocity of replication forks (kb/min)** | | |
|  | Number of values | 205 | 127 |
|  | Minimum | 0.6995 | 0.5525 |
|  | 25% Percentiles | 1.528 | 2.13 |
|  | Median | 1.724 | 2.408 |
|  | 75% Percentiles | 1.935 | 3.01 |
|  | Maximum | 3.212 | 4.053 |
|  | Mean | 1.754 | 2.56 |
| **Asymmetry of replication forks (long/short fork ratio)** | | | |
| Number of values | | 64 | 39 |
| Minimum | | 1 | 1 |
| 25% Percentiles | | 1.02 | 1.012 |
| Median | | 1.068 | 1.097 |
| 75% Percentiles | | 1.226 | 1.199 |
| Maximum | | 2.184 | 2.645 |
| Mean | | 1.163 | 1.189 |
| **Lengths of the analysed fibres (kb)** | | | |
| Number of values | | 187 | 151 |
| Minimum | | 104.3 | 171.4 |
| 25% Percentiles | | 330.1 | 361.2 |
| Median | | 446.9 | 446.9 |
| 75% Percentiles | | 516.9 | 511.6 |
| Maximum | | 1363 | 1146 |
| Mean | | 459.6 | 464.3 |
| **Estimated number of active origins per cell** | | | |
| Genomic sequence (50.081 Mb): median IOD | | 329 | 235 |
| Genomic sequence (50.081 Mb): mean IOD | | 279 | 228 |

**DNA replication parameters obtained by DNA combing in two cell types (PCF and BSF) of *T. brucei***. The estimated numbers of origins were obtained by dividing the genomic sequence length (50.081 Mb) by either the median or the mean IODs calculated on combed fibres.

#### SUPLEMENTARY FIGURES

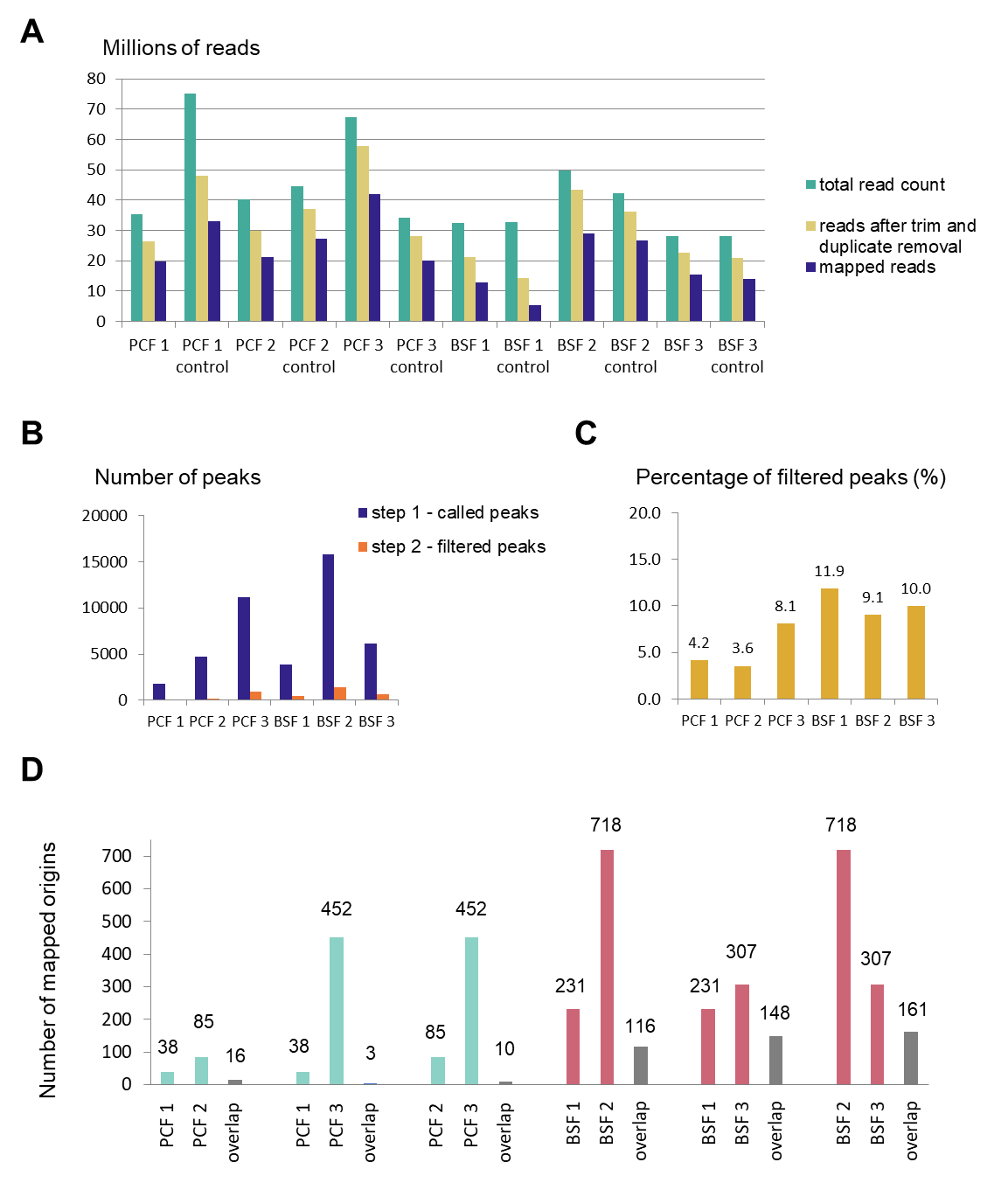

##### Supplementary Figure 1

A) Number of reads obtained by paired-end sequencing (starting number), after trimming and duplicate removal and after mapping of reads against *T. brucei* 427-2018 genome. B) Number of peaks after peak calling step (bars in blue) and after peak filtering step (bars in orange). Three replicates of PCF and BSF cells are presented. C) Proportion of the filtered peaks compared to the called peaks (100%). Three replicates of PCF and BSF cells are presented. D) Overlap of mapped origins between three replicates of PCF and BSF cells (±100 bp window). The number of mapped origins in three replicates of PCF and BSF cells and their overlap are shown above the bars. The overlap is represented by grey bars.

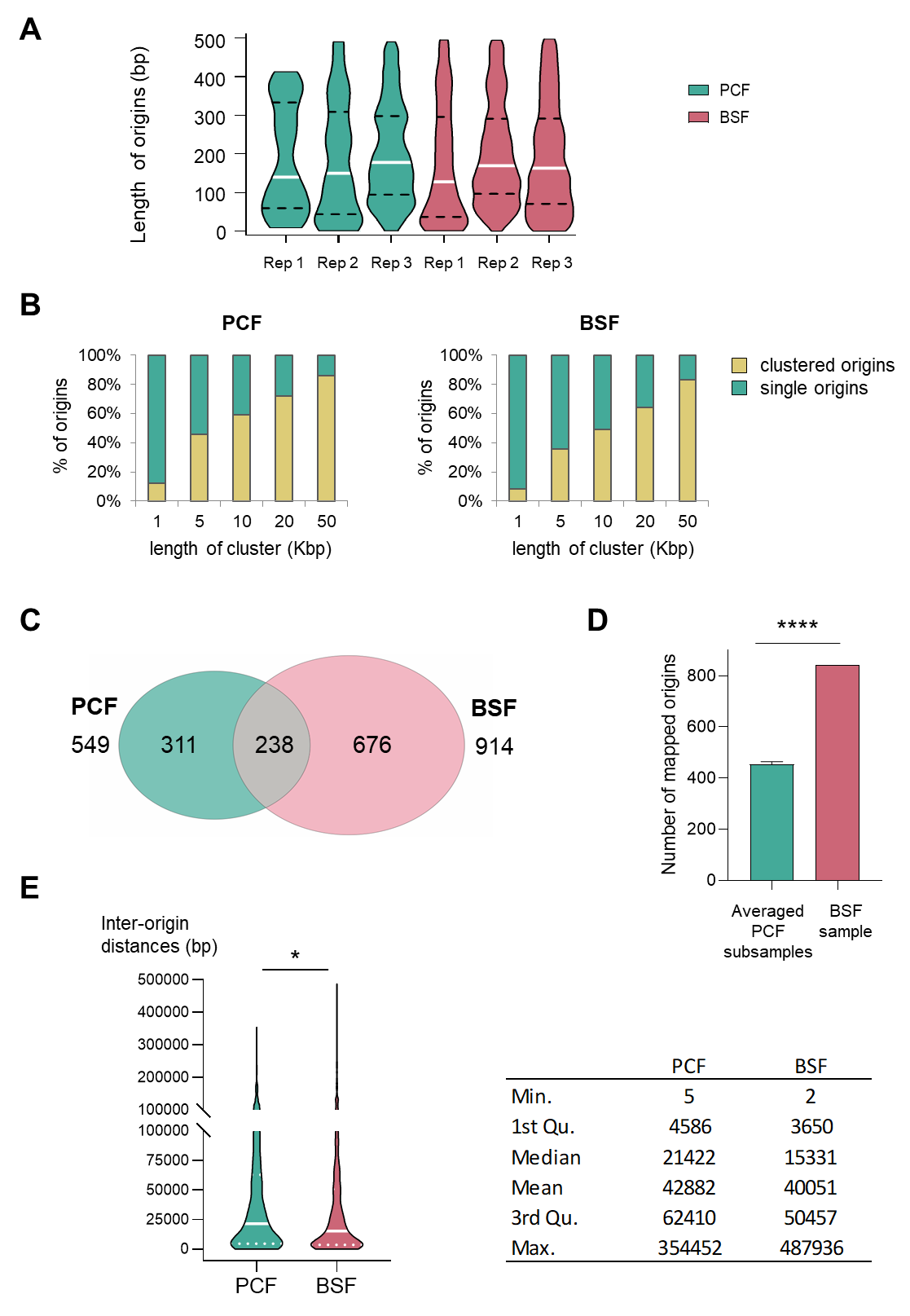

##### Supplementary Figure 2

A) Violin-plots present the distribution of the lengths of mapped origins for the three replicates of PCF and BSF cells. The white line represents the median value. The dotted lines represent the 25^th^ (Q1) and 75^th^ (Q3) percentiles of the measurements. B) The graphs present the proportions of clustered and single origins within genomic regions of indicated lengths for PCF (left) and BSF (right) cell lines. C) Venn diagram showing the number of mapped origins in PCF and BSF cells, with the number of common origins for both cell types. Direct overlap was calculated. D) Bar plots of the normalized numbers of detected origins in PCF (green) compared to BSF (pink). For the normalization all reads of the three replicates of BSF and PCF were combined respectively and the number of reads in PCF was down-sampled 15 times so that the number of mapped reads match the one in BSF. Afterwards strand-specific peak-calling against normalized SNS-depleted control was performed, followed by peak filtering and origin detection, as described in the Methods and in the main text. The mean for the 15 down sampled PCF subsamples was calculated; the error bars represent the standard deviation. The *P*-value was calculated using a t-test (**** - *P*<0.0001). E) Distribution of population based inter-origin distances of PCF and BSF cells represented as violin plots, green for PCF cells and pink for BSF cells. The white line represents the median value. The dotted lines present the 25^th^ (Q1) and 75^th^ (Q3) percentiles of measurements. A two-tailed Mann-Whitney test was used to compute the *P*-value (* - *P*<0.05). The table shows the minimum (Min.) and maximum (Max.) value, the 25^th^ (1st Qu.) and 75^th^ (3rd Qu.) percentiles as well as the median and mean values. The measurements are indicated in base pairs.

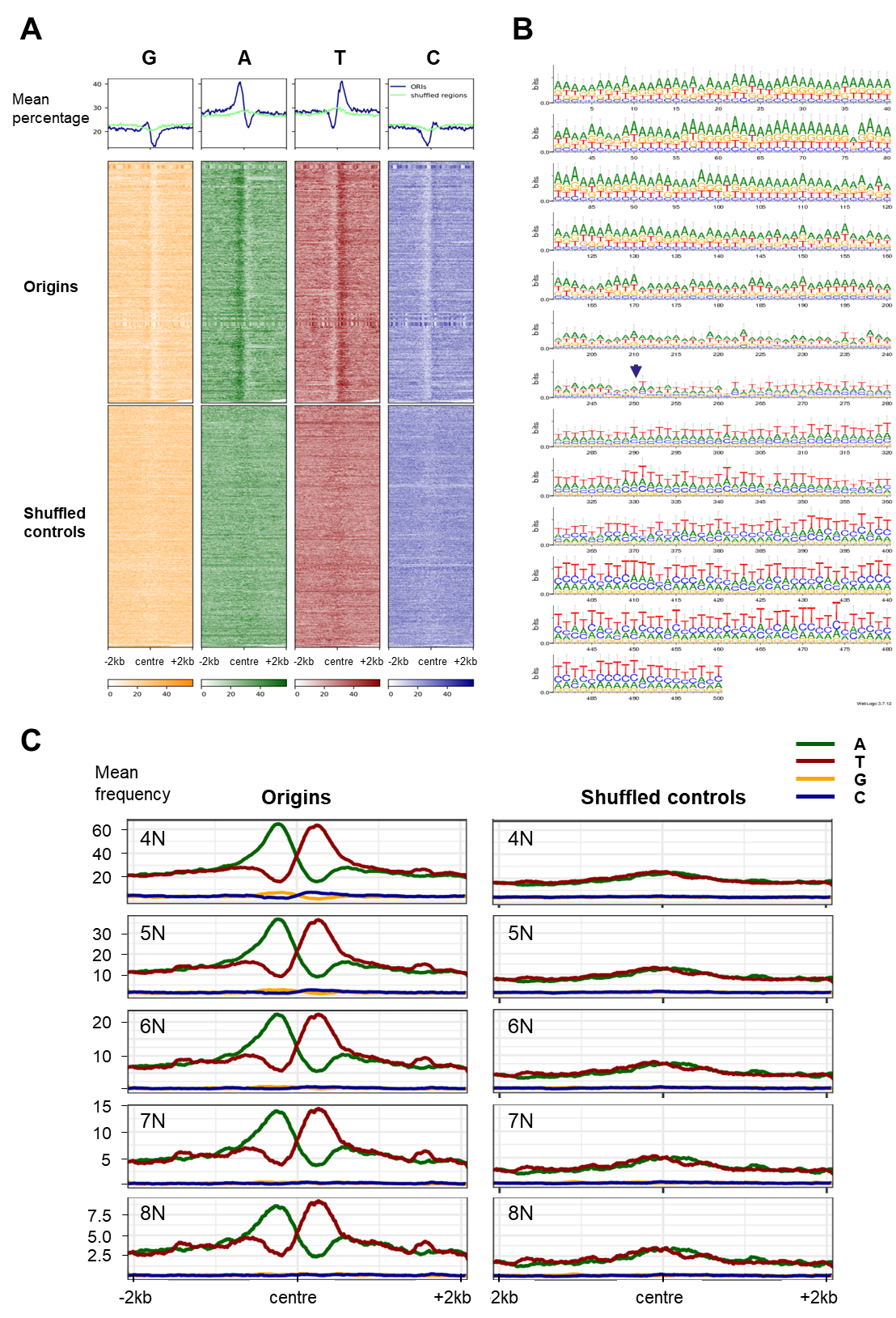

##### Supplementary Figure 3

A) Profile plots and heatmaps presenting the distribution of the four nucleotides around centred origins (blue line in the profile plot) and shuffled controls (green line in the profile plot). A region of ± 2 kb from the centre was analysed. The nucleotide percentage was calculated within 20 bp windows. B) A graphical representation of nucleotide occurrence. Sequence logo of the centred origins (± 250 nt) generated with WebLogo 3 ^47^ (<https://weblogo.threeplusone.com/>). The arrow indicates the centre of the averaged origins. C) Profile plots showing polynucleotide frequency around centred origin (left panels) and shuffled controls (right panels). A region of ± 2 kb from the centre was analysed. The frequency of four to eight polynucleotides (4N-8N) is shown as indicated.

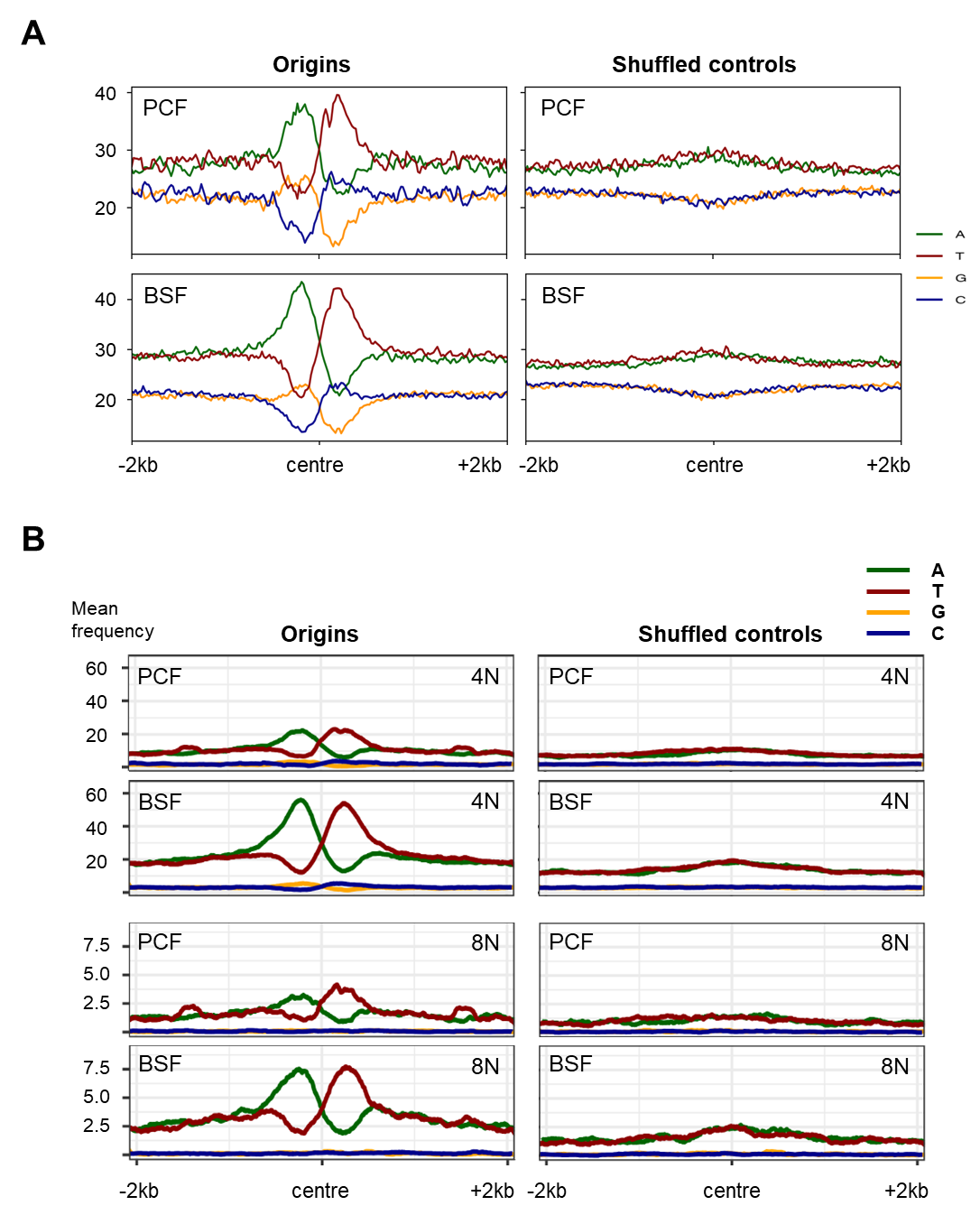

##### Supplementary Figure 4

A) The profile plots illustrate the distribution of four nucleotides around centred origins from PCF and BSF cells separately and corresponding shuffled controls (± 2 kb). The nucleotide percentage was calculated within 20 bp windows. The mapped origins in the PCF and BSF cells were analysed separately to show that they can be compared with the merged (PCF+BSF) set of origins shown in the main figures. B) Profile plots showing polynucleotide frequency around centred origin from PCF and BSF cells (left panels) and corresponding shuffled controls (right panels). A region of ± 2 kb from the centre was analysed. The frequency of four and eight polynucleotides (4N and 8N) is shown as indicated. The mapped origins in the PCF and BSF cells were analysed separately to show that they can be compared with the merged (PCF+BSF) set of origins shown in the main figures.

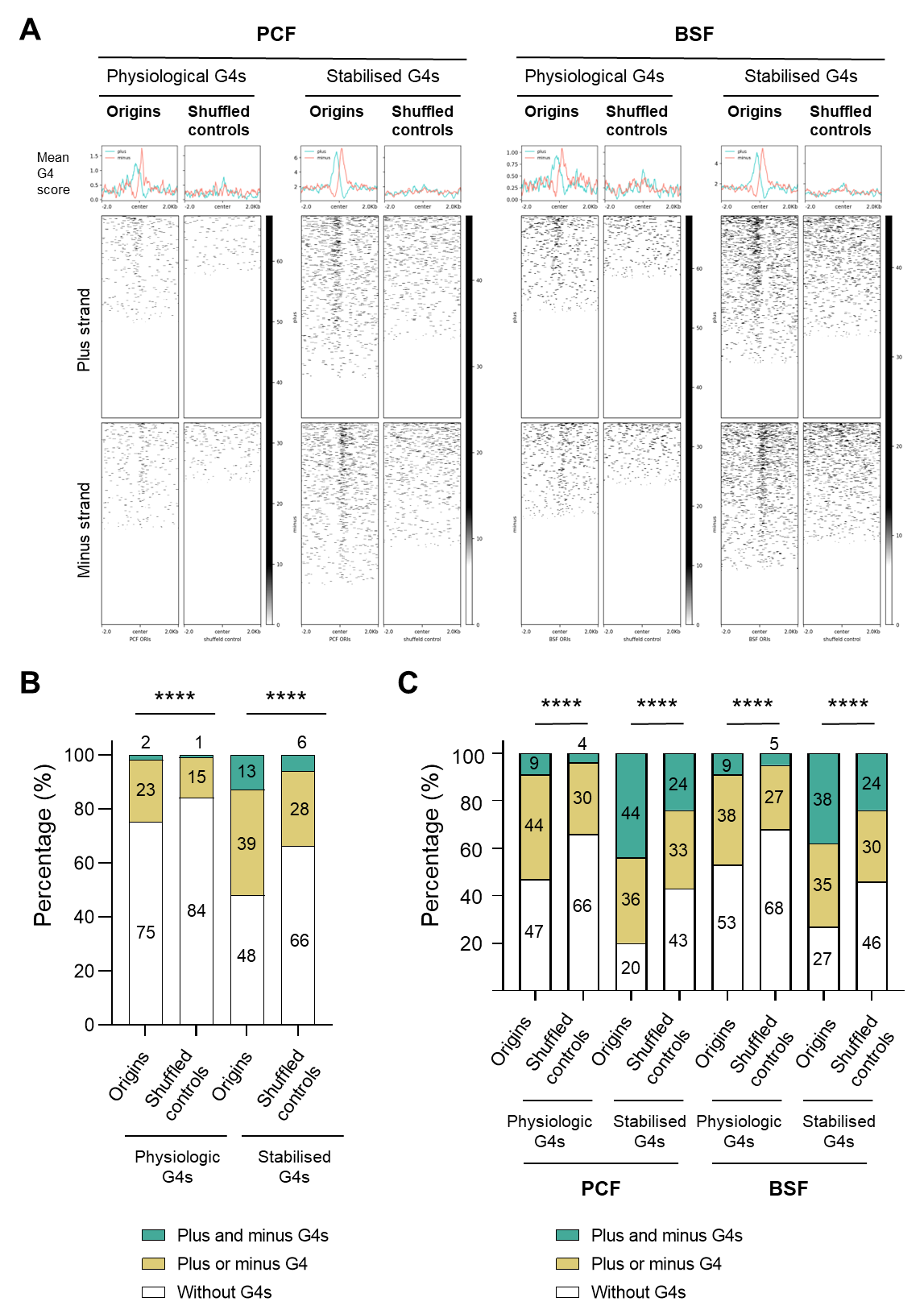

##### Supplementary Figure 5

A) The profile plots and heat maps show the distribution of the experimentally obtained G4 structures around centred origins and intergenic shuffled controls (± 2 kb) in the Tbb TREU927 reference genome. The mapped origins in the PCF and BSF cells were analysed separately to show that they can be compared with the merged (PCF+BSF) set of origins shown in the main figures. The plus strand (light blue) and minus strand G4s (pink), obtained under physiological conditions and in the PDS drug stabilized condition ^49^ were overlapped with the stranded SNS-seq mapped origins. Mean G4 score presents average G4 value ^49^ per 20 bp window. B) The proportions of origins (merged PCF+BSF) and corresponding shuffled control regions that lack or possess G4 structures on one or both sides of the centre are shown in percentage. A ±0.5 kb window from the centre was analysed. Two sets of experimental G4 structures (physiologic G4s and stabilised G4s) ^48^ were analysed in origins and intergenic shuffled control. The *P*-values were calculated using the Chi-square test, which involved a comparison of the absolute numbers of three indicated categories for each pair of datasets (**** - *P*<0.0001). C) The proportions of origins that lack or possess G4 structures ^49^ on one or both sides of the centre were determined. The mapped origins in the PCF and BSF cells were analysed separately to show that they can be compared with the merged (PCF+BSF) set of origins shown in the main figures. The ± 2 kb region from the centre was analysed for the presence of G4s. Two sets of experimental G4 structures, physiological and stabilised ^49^ were subjected to analysis in comparison with an intergenic shuffled control. The P-values were calculated using the Chi-square test, which involved a comparison of the absolute numbers of indicated categories for each pair of datasets (**** - P<0.0001).

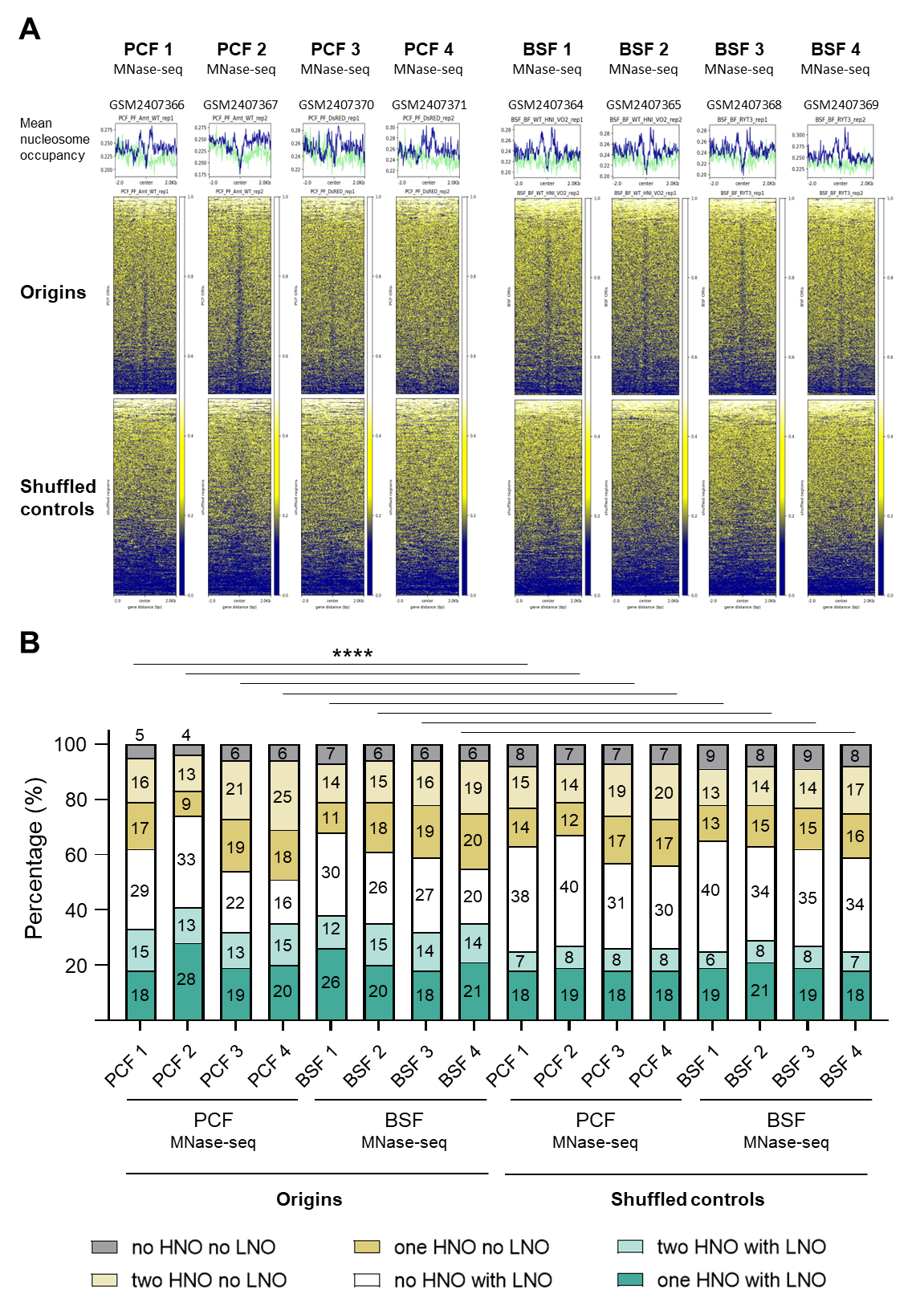

##### Supplementary Figure 6

A) Profile plots and heatmaps presenting the distribution of nucleosomes around the centred origins and shuffled controls. The region of ± 2 kb from the centre was analysed. Eight replicates of MNase-seq data (Four PCF replicates: GSM2407366, GSM2407367, GSM2407370, GSM2407371 and four BSF replicates: GSM2407364, GSM2407365, GSM2407368 and GSM2407369) describing the distribution of nucleosomes ^50^ were analysed. The upper profile plots depict the origins and shuffled controls, which were represented in blue and green, respectively. Mean nucleosome occupancy presents average nucleosome score (dyad values) ^50^ per 20 bp window. B) The proportions of origins and shuffled controls with or without high nucleosome occupancy (HNO) region on one or both sides of the centre and a low nucleosome occupancy (LNO) at the centre of the regions in different combinations are shown in stacked bar plots. The proportions were calculated for the eight replicates of nucleosome positioning as indicated ^50^ . The *P*-values were calculated using the Chi-square test, which involved a comparison of the absolute numbers of six indicated categories for each pair of datasets (**** - *P*<0.0001).

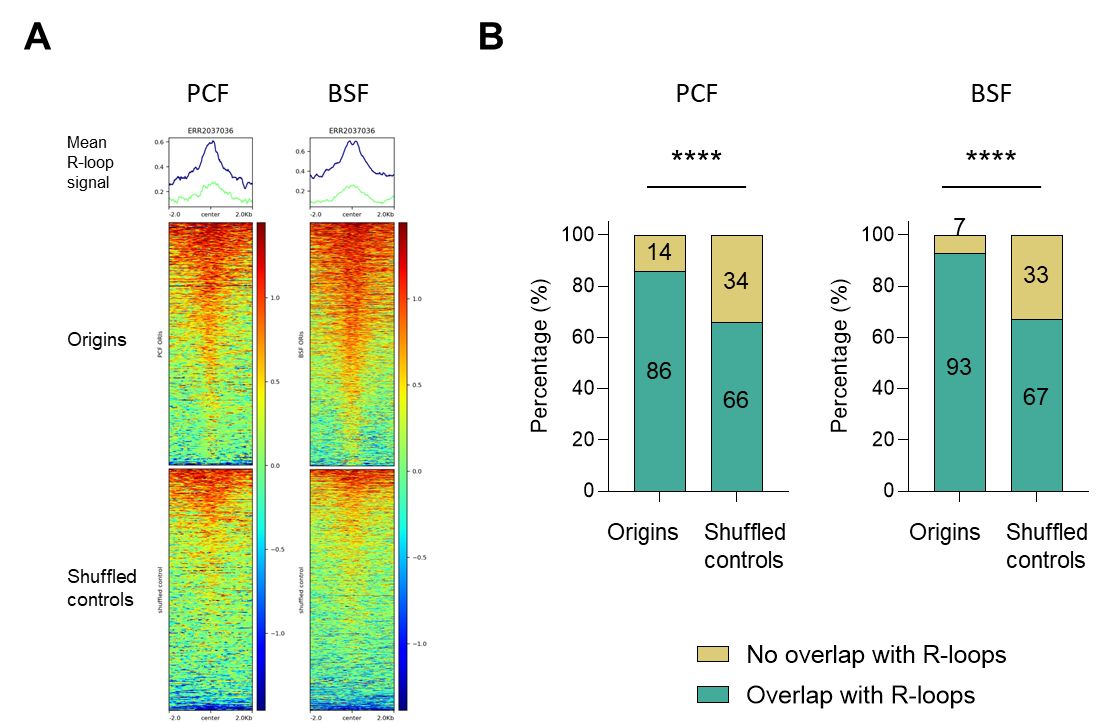

##### Supplementary Figure 7

A) The profile plots and heatmaps show the distribution of RNA:DNA hybrids (R-loops) ^54^, around the centred origins of PCF and BSF cells (blue line) and the corresponding intergenic shuffled controls (green line) (± 2 kb). Mean DRIP-seq signal presents average DRIP-seq signal ^54^ per 20 bp window. B) The percentage of origins of PCF and BSF cells and corresponding shuffled controls that overlap with R-loops (overlap calculated without window). The *P*-values were calculated using Fisher's exact (two-sided) test, which involved a comparison of the numbers of indicated categories for each pair of datasets (**** - *P*<0.0001). The mapped origins in the PCF and BSF cells were analysed separately to show that they can be compared with the merged (PCF+BSF) set of origins presented in the Figure 7 and in the main text.

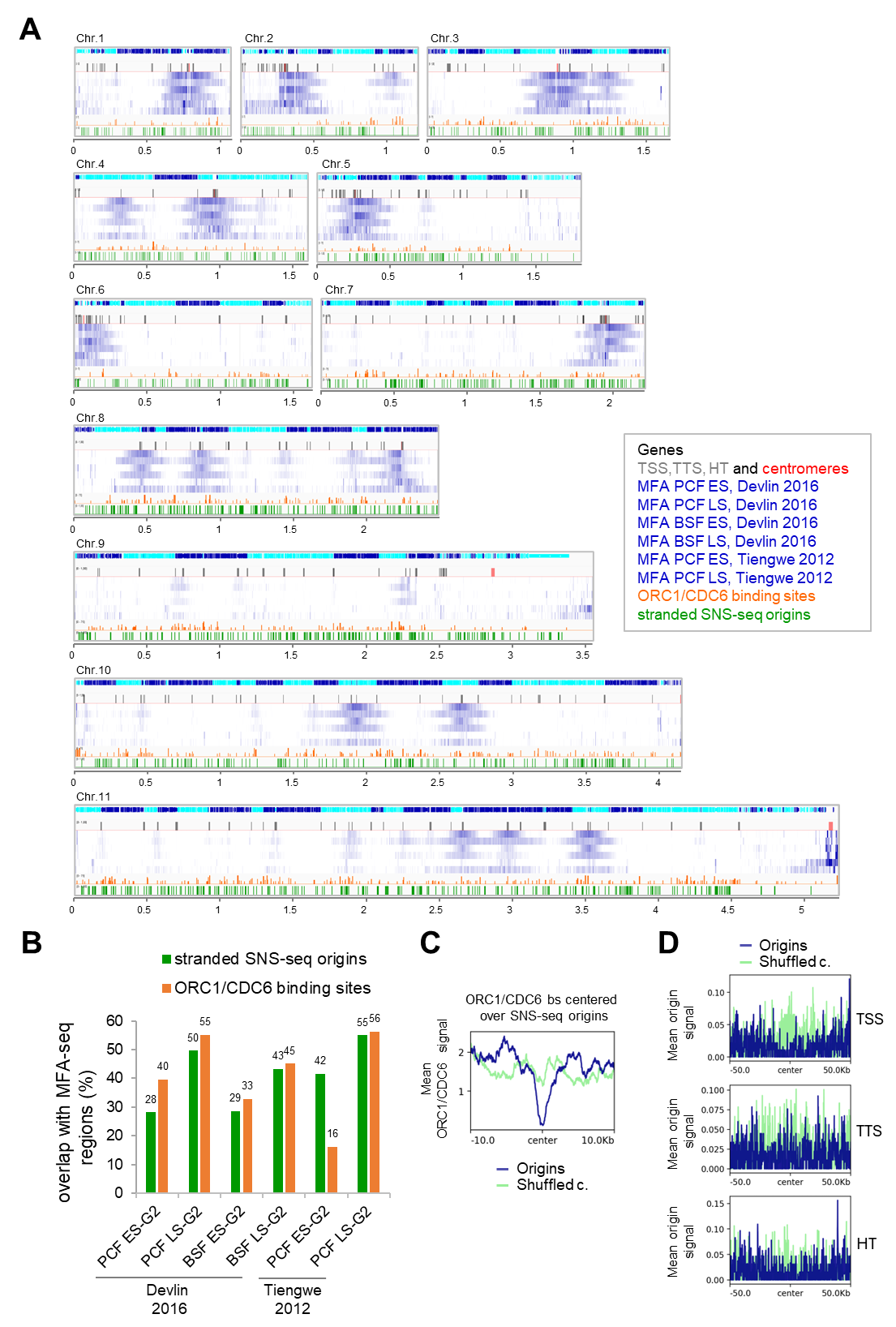

##### Supplementary Figure 8

A) Chromosomal overview of MFA-seq and stranded SNS-seq origins from Integrative Genomics Viewer (IGV) (scale in Mb). The 11 core chromosomes of the Tb Lister 427-2018 reference genome are shown. The first track shows genes and their orientation (blue presents genes on the plus strand and turquoise presents genes on the minus strand). The second track shows centromeres in red (for chromosomes 9-11 the centromeres are not mapped to the core chromosome) and in grey TSS, TTS and HT regions ^56^. Tracks 3-8 show MFA-seq data ^42, 43^ displaying the ratio of read depth between G2 and S phase in blue (scale 1-1.8). Tracks 3 and 4 show MFA-seq data from PCF cells (track 3 - early S phase to G2 phase ratio and track 4 - late S phase to G2 phase ratio). Tracks 5 and 6 show MFA-seq data from BSF cells (track 5 - early S phase to G2 phase ratio and track 6 - late S phase to G2 phase ratio) ^43^. Tracks 7 and 8 show MFA tracks from PCF cells. (Track 7 represents the early S phase to G2 phase ratio and track 8 represents the late S phase to G2 phase ratio) ^42^ (data obtained from Richard McCulloch via personal communication). Track 9 shows ORC1/CDC6 binding sites (retrieved from TriTrypDB) in orange. Track 10 shows the positions of stranded SNS-seq origins in green. B) Bar plot presenting the percentage of the overlap of stranded SNS-seq origins (in green) and ORC1/CDC6 binding sites (in orange) with MFA-seq regions from the indicated data sets. Overlap was calculated without a window of 100 nt. The first 4 bars are overlap from Devlin *et al*. (2016) ^43^ data; the last 2 bars are overlap from Tiengwe *et al*. (2012) ^42^ data. Overlap of stranded SNS-seq origins with early S phase to G2 phase MFA data sets are shown in light blue and overlap with late S phase to G2 phase MFA data sets are shown in dark blue. C) Profile plots illustrating the distribution of Orc1/Cdc6 binding sites ^42^ around centred origins (± 2 kb upper panel and ± 10 kb lower panel) for merged origins (PCF and BSF) in blue and corresponding shuffled control in light green. The mean Orc1/Cdc6 signal was calculated within 20 bp windows. D) Profile plots illustrating the distribution of merged origins (PCF and BSF) in blue and corresponding shuffled control in light green around centred TSS (upper panel), TTS (middle panel) and HT regions (lower panel) ^56^ (± 50 kb). The mean origin signal was calculated within 20 bp windows.

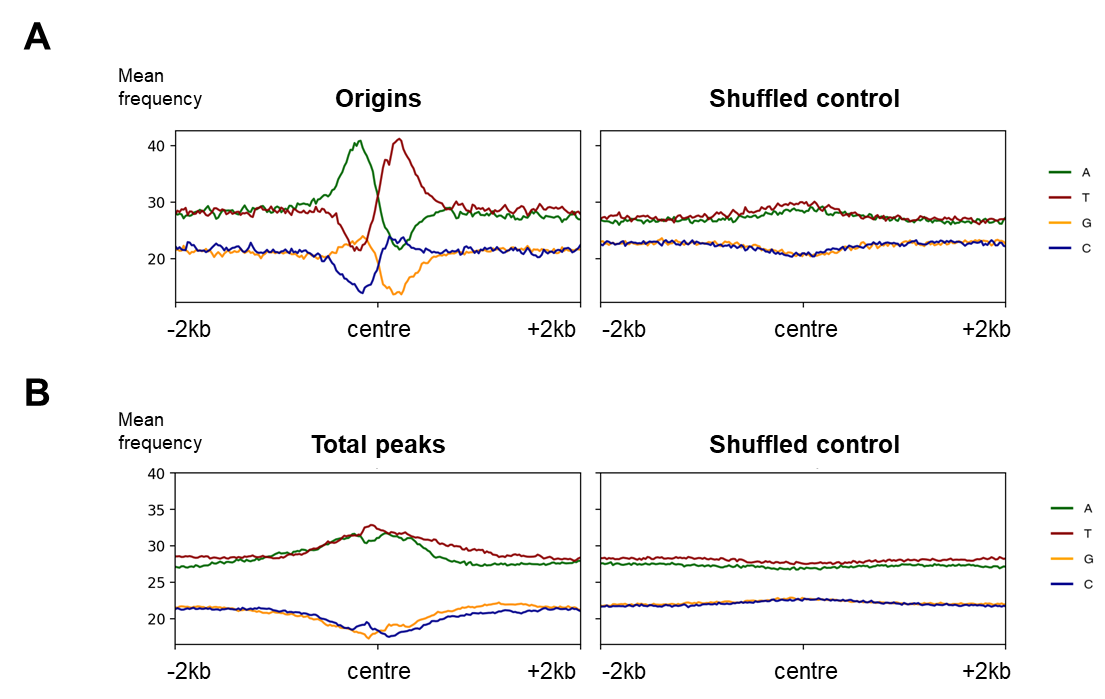

##### Supplementary Figure 9

The profile plots show nucleotide distributions around two sets of regions derived from the same stranded SNS-seq experiments. A) The merged origins (PCF + BSF), mapped with stranded SNS-seq approach in the middle of two divergent SNS peaks (see Methods), and their corresponding shuffled controls (± 2 kb) are shown. B) Total called peaks, which would represent origins in a conventional (non-stranded) SNS-seq approach, along with their respective shuffled controls (± 2 kb) are shown. The nucleotide percentages were calculated in 20 bp windows.

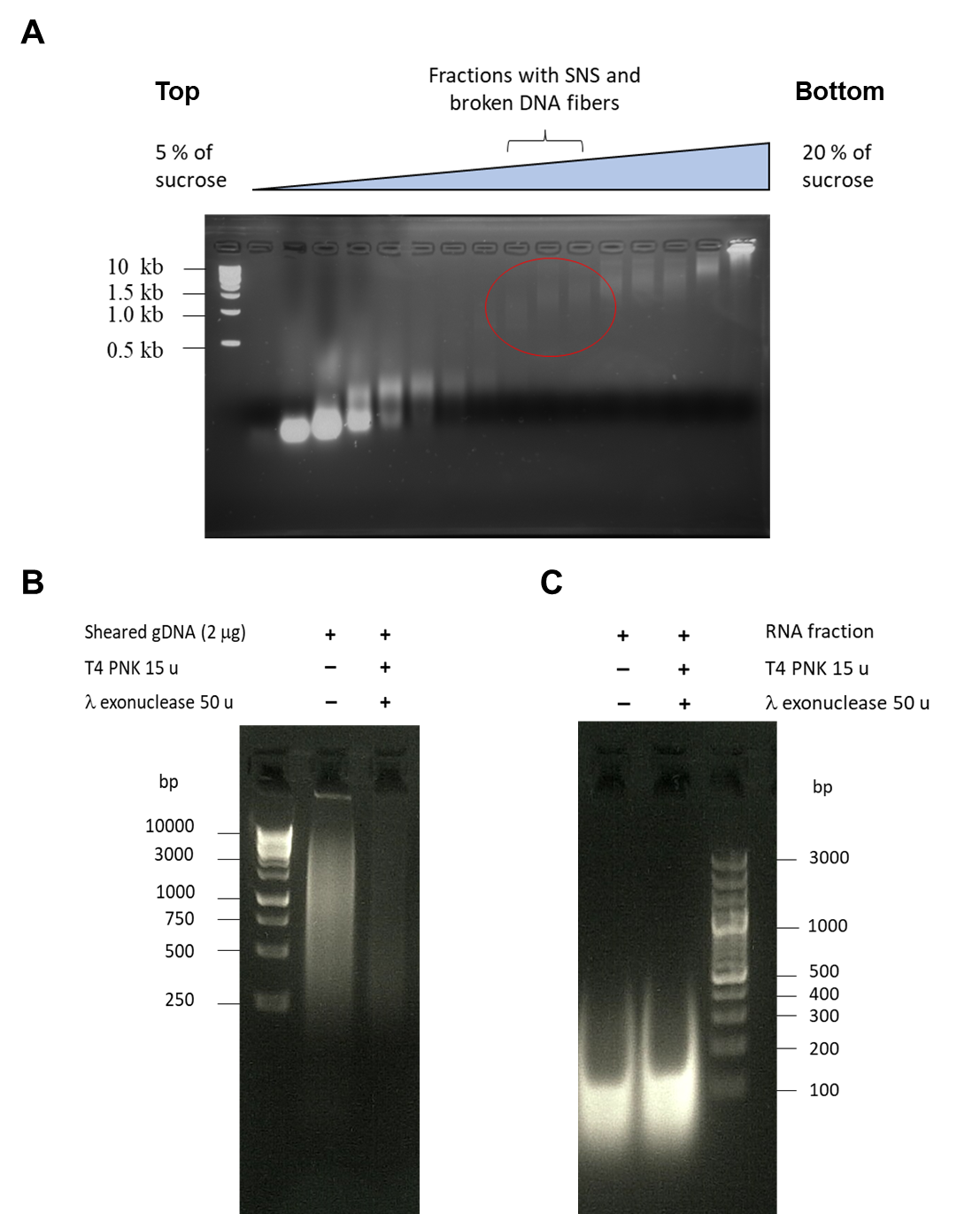

##### Supplementary Figure 10

A) Native agarose gel electrophoresis of fractions from sucrose gradients. gDNA was denatured, centrifuged on 5-20 % sucrose gradient and fractions were collected from the top (5% sucrose) to the bottom of gradient (20% sucrose). The size of DNA molecular marker is indicated on the right. Fractions containing nucleic acids of 0.5 to 2.5 kb are encircled. B) Control of the efficiency of the T4 PNK and λ-exonuclease enzymes. gDNA was fragmented to the size of SNS fibres and 2 µg of starting gDNA was treated once with the indicated number of units of T4 PNK and λ-exonuclease enzymes to check the efficiency of enzymes. The SNS containing fractions were treated under the same condition but 3-4 times. The size of the DNA molecular marker is indicated. C) RNA is preserved during treatment with T4 PNK and λ-exonuclease enzymes. RNA from the top fractions of the sucrose gradient was treated with the indicated number of units of T4 PNK and λ-exonuclease enzymes to check if the RNA fibres are preserved after this treatment. The size of the DNA molecular marker is indicated.
